# Nociceptor clock genes control excitability and pain perception in a sex and time-dependent manner

**DOI:** 10.1101/2025.04.07.646998

**Authors:** Aurélie Brécier, Courtney A. Bannerman, Yu-Feng Xie, Christopher Dedek, Amanda M. Zacharias, Ciara D. O’Connor, Steven D. Miller, Laurel L. Ballantyne, Justin Du Bois, Qingling Duan, Steven A. Prescott, Nader Ghasemlou

## Abstract

Nociception is critical for pain perception and survival and begins with the activation of nociceptors, specialized sensory neurons located in the dorsal root ganglia (DRGs). Both sex and circadian rhythms, governed by clock genes, seem to play a significant role in modulating pain perception. However, the potential interaction between circadian rhythms and sex differences in nociception at the peripheral level has been largely overlooked. Here, we first report that DRGs from mice express clock genes in a time- and sex-dependent manner. Using whole-cell recordings in whole-mounted DRGs and optogenetic stimulation of Nav1.8-expressing neurons, we demonstrate that male nociceptors exhibit reduced excitability during the night, while female nociceptor excitability remains stable across time points. Disruption of the core clock gene *Bmal1* in Nav1.8-expressing neurons not only diminished nociceptor activity but also abolished the nighttime reduction in heat sensitivity, highlighting a pivotal role for the molecular clock in regulating nociception. Transcriptomic analyses, voltage-clamp recordings, and pharmacological experiments identified the voltage-gated chloride channel ClC-2, controlled by *Bmal1*, as a key mediator for the observed fluctuations in male nociceptor excitability. This work opens new avenues for chronobiology-inspired strategies in pain management tailored to sex-specific mechanisms.

## INTRODUCTION

Males and females not only process and experience pain differently, but the molecular and cellular mechanisms underlying these effects are often sexually dimorphic ^1–4^. Much of this work has focused on chronic pain, with the activation of pathways in the central and peripheral nervous systems playing important roles in this response. Less well-studied are changes in basal nociception, the neural process of encoding and processing noxious stimuli that are critical for pain perception and survival. Activation of the specialized sensory neurons that reside in the dorsal root ganglia (DRGs), called nociceptors, results in the transduction of harmful thermal, mechanical, and chemical stimuli into electrical signals transmitted to the central nervous system. The functional properties of nociceptors, particularly their excitability, directly influence the perception of pain ^5^.

Circadian rhythmicity and the endogenous clock act on most physiological systems, providing a unique opportunity to dissect potential mechanisms regulating pain sensitivity ^6,7^; targeting this rhythmicity may help inform novel and/or time-specific therapeutic strategies for pain management. In humans, circadian patterns were reported in pain sensitivity with lower sensitivity during specific phases of the day ^7–10^. Importantly, the work by Daguet and colleagues showed that this change in baseline sensitivity, assessed in males only, was due to endogenous circadian rhythms and not sleep. Similar rhythms have been documented in rodents, where studies demonstrated diurnal variations in pain thresholds ^11–17^, though differences between males and females have been rarely assessed ^18^. The potential interaction between circadian rhythms and sex differences in nociception has been largely overlooked, and the role of the peripheral system in this phenomenon is completely unknown. While there is evidence that DRGs possess a local transcriptional clock ^19–22^, it is unclear whether these rhythms can impact nociceptor excitability and pain perception.

Understanding the intersection of circadian rhythms, sexual dimorphism of pain, and nociceptor function is essential for elucidating the peripheral contributions to rhythmic pain regulation. Here, we report i) a sex-dependent diurnal variation in nociceptor excitability using *ex vivo* recordings and optogenetic behavioral testing; ii) a pivotal role for clock genes in controlling the daily fluctuation of excitability and pain thresholds following the specific loss of the molecular clock in nociceptors; and iii) the critical role of the voltage-gated chloride channel ClC-2 in the circadian fluctuation of pain in male but not female mice using both transcriptomic analysis and pharmacological targeting of the rhythmically expressed channel in nociceptors *ex vivo* and *in vivo*. This study lays the groundwork for a better understanding of variations in nociceptor excitability between sexes and the impact of circadian rhythms and may provide new avenues of study for therapeutic opportunities to treat acute and chronic pain.

## RESULTS

### DRGs express clock genes in a time and sex-dependent manner

Previous work has shown that DRGs express the main clock genes in a circadian manner ^21,22^. However, it is unclear if the clock machinery is comparable between males and females, and whether this could explain the fluctuations observed in nociception. We therefore collected DRGs from male and female mice across four times of day in 12h light:12h dark conditions. Under such lighting conditions, Zeitgeber time (ZT) is used with ZT0 denoting lights on, and ZT12 lights off in the facility. Here, whole DRGs were collected at ZT2, ZT8, ZT14, and ZT20 (n = 3 mice/time point/sex) to compare the expression of core clock genes (Fig. 1a-h). We found similar changes across sexes in the rhythmic expression of the master clock gene *Bmal1*, as well as for *Nr1d1* and *Per2* (R^2^ ≥ 0.65); *Per1* expression also fit to a 24 h sine wave in both sexes, though less well (R^2^ = 0.37-0.40). Meanwhile, expression of *Clock*, *Cry1*, and *Cry2* remained stable across the light/dark cycle in both male and female DRGs. The expression of *Nr1d2* also followed a diurnal rhythm in females but not in males (Fig. 1d). Of note, *Per1* transcript levels were significantly different in female DRGs than in males (Fig. 1e). Overall, these results are largely in line with previous findings ^21,22^ and taken together suggest that the molecular clock in the DRG of male and female mice is largely similar, but with some notable differences. These differences may reflect variations in gene expression, regulatory mechanisms, or interactions with hormonal or environmental factors. Although the core components of the clock appear to be conserved, the subtle differences could influence rhythmic regulation of nociceptive processing, potentially contributing to sex-specific variations in pain sensitivity and chronobiology.

**Fig. 1.**
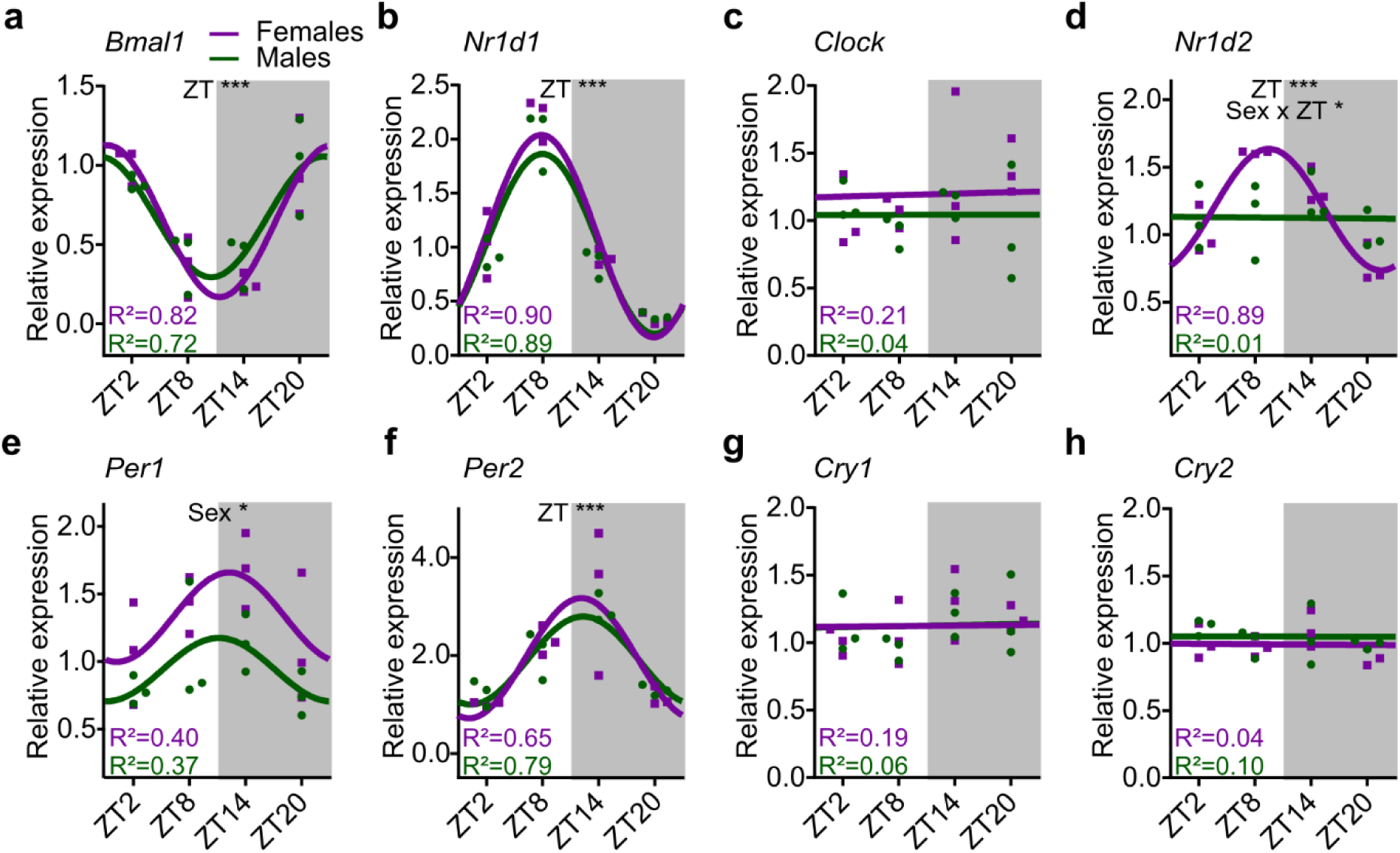
Sex difference in DRG clock-gene expression. **a-h**. Best fitted model (24h sines wave or line) of *Bmal1, Nr1d1, Clock, Nr1d2, Per1, Per2, Cry1* and *Cry2* expression in female (purple) vs in male (green) DRGs. Extra sum-of-squares F tests were performed with R² values representing the degree of fitting. Data were normalized to ZT2 in females (n = 12 mice per sex). **a.** Two-way ANOVA with repeated measures, sex effect F (1, 4) = 0.049, P = 0.83, ZT effect F (1.2, 5.0) = 40, P = 0.001, sex x ZT interaction F (3, 12) = 1.0, P = 0.42. **b.** Two-way ANOVA with repeated measures, sex effect F (1, 4) = 0.88, P = 0.40, ZT effect F (2.0, 7.9) = 104, P < 0.001, sex x ZT interaction F (3, 12) = 0.36, P = 0.78. **c.** Two-way ANOVA with repeated measures, sex effect F (1, 4) = 1.3, P = 0.33, ZT effect F (1.9, 7.6) = 0.83, P = 0.47, sex x ZT interaction F (3, 12) = 1.1, P = 0.39. **d.** Two-way ANOVA with repeated measures, sex effect F (1, 4) = 0.77, P = 0.43, ZT effect F (1.5, 6.1) = 7.9, P = 0.024, sex x ZT interaction F (3, 12) = 4.0, P = 0.035. **e.** Two-way ANOVA with repeated measures, sex effect F (1, 4) = 11, P = 0.031, ZT effect F (1.7, 6.6) = 3.3, P = 0.11, sex x ZT interaction F (3, 12) = 0.18, P = 0.91. **f.** Two-way ANOVA with repeated measures, sex effect F (1, 4) = 0.031, P = 0.87, ZT effect F (1.4, 5.6) = 16, P = 0.006, sex x ZT interaction F (3, 12) = 0.35, P = 0.79. **g.** Two-way ANOVA with repeated measures, sex effect F (1, 4) = 0.059, P = 0.82, ZT effect F (1.7, 6.9) = 1.7, P = 0.25, sex x ZT interaction F (3, 12) = 0.32, P = 0.81. **h.** Two-way ANOVA with repeated measures, sex effect F (1, 4) = 1.2, P = 0.33, ZT effect F (1.9, 7.5) = 1.5, P = 0.29, sex x ZT interaction F (3, 12) = 0.54, P = 0.66. * P < 0.05, ** P < 0.01, *** P < 0.001.

### Excitability of male but not female nociceptors is decreased during the dark phase

Since the DRG consists of a significant proportion of satellite glial cells and non-nociceptive sensory neurons, the transcriptomic changes observed do not provide definitive evidence that nociceptor activity is regulated by circadian (or diurnal) rhythms. In addition, transcriptomic changes, even in nociceptors, do not necessarily convey any functional changes. We therefore used electrophysiology to test if nociceptors experience functional changes that might derive from transcriptional changes. We examined the excitability of single sensory neurons at ZT2 and ZT14, corresponding to peak and trough expression of most circadian genes in both male and female DRGs. To do so, we performed whole-cell recordings on freshly harvested L3/4 DRG at the specified time points following the guidelines of a published method ^23^ (Fig. 2a-b). Small-diameter neurons (< 25µm) were targeted to specifically record putative nociceptors (Fig. 2c). We observed a sex effect in the resting membrane potential and rheobase of nociceptors, with a higher resting membrane potential and a lower rheobase measured in nociceptors from female mice (Fig. 2d-f). We found no difference in membrane capacitance or resistance between males and females, nor between ZT2 (light phase) and ZT14 (dark phase; Extended Data Fig. 1a-b). These results confirm previous findings performed *in vivo* and *in vitro* ^24,25^ and could explain the sex difference observed at the behavioral level ^1^. In addition, we noticed a significant decrease in the number of APs evoked by pulses > 200pA at ZT14 compared to ZT2 in nociceptors from male DRG but not from female DRG (Fig. 2g). Of note, the evoked activity is similar between males and females at ZT2 but significantly differs at ZT14 (Extended Data Fig. 1c-d), suggesting a sex-specific diurnal regulation of nociceptor excitability. To investigate the sensory implications of this variation in nociceptor excitability, we optogenetically stimulated the hind paw of mice selectively expressing channelrhodopsin-2 (ChR2) in Nav1.8+ neurons (Fig. 2h). Our results indicate that withdrawal latency remains constant in female mice but was longer during the dark phase in male mice (Fig. 2i), consistent with the male-specific drop in excitability at ZT14. Overall, these data suggest that the excitability of Nav1.8-expressing neurons differs in a sex- and time-dependent manner, as observed in both *ex vivo* and *in vivo* conditions. Thus, the intrinsic properties of Nav1.8+ sensory neurons appear to be modulated by diurnal and sex-specific regulatory mechanisms.

**Fig. 2.**
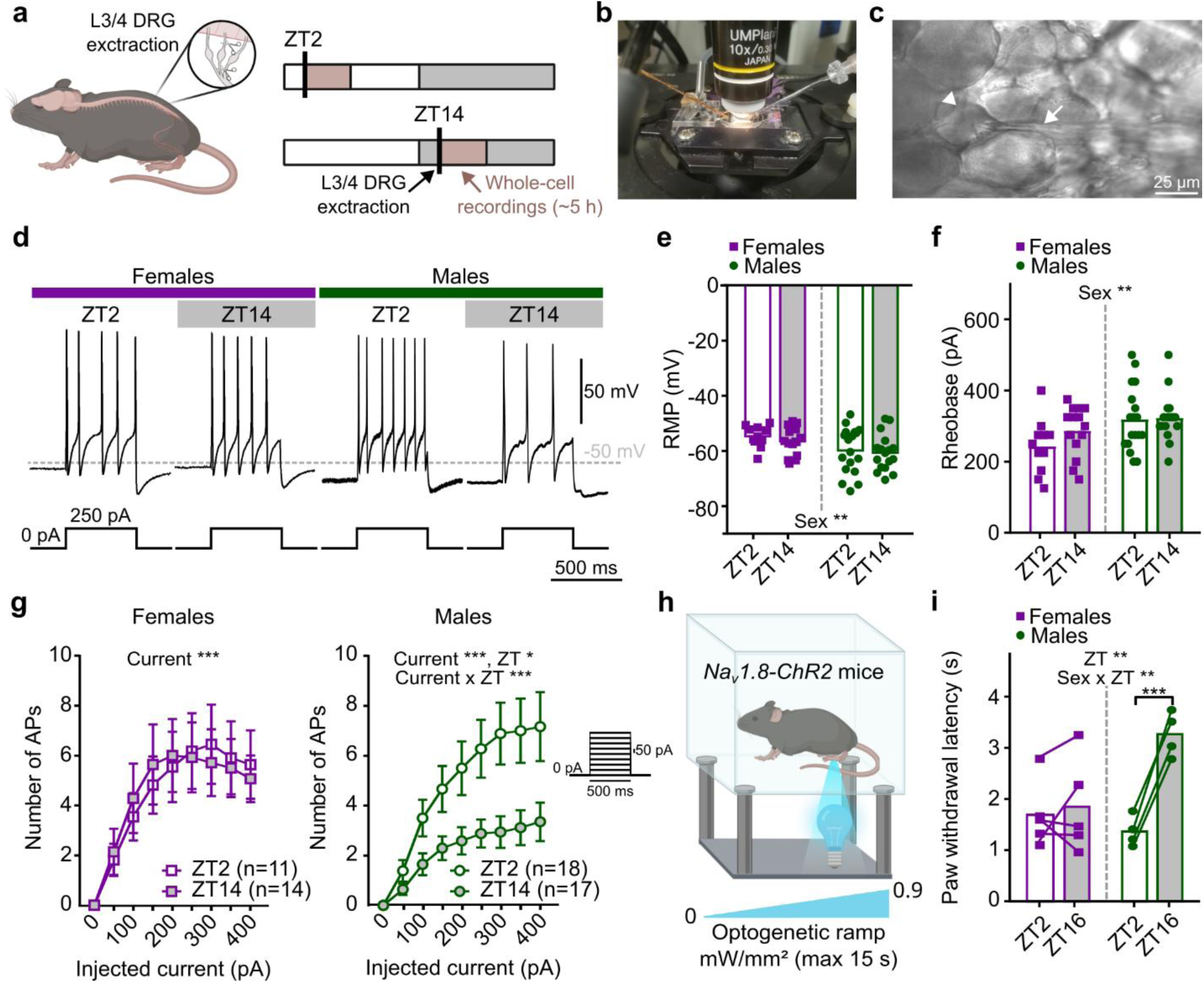
Nociceptor excitability differs between female and male mice, and day vs night. **a.** Protocol for intact DRG whole cell recordings at ZT2 and ZT14. **b**. Experimental set-up for whole-cell recordings of freshly harvested DRGs. **c.** Small diameter DRG neuron (arrow head) recorded in whole cell mode via a glass pipette (small arrow). **d.** Activity evoked by 250 pA injected for 500 ms in female and male mice nociceptors at ZT2 and ZT14. **e.** Resting membrane potential. Mixed-effects model, sex effect F (1, 56) = 7.4, P = 0.009, ZT effect F (1, 56) = 0.63, P = 0.43, sex x ZT interaction F (1, 56) = 0.073, P = 0.79, n > 5 mice per group, bars represent mean. **f.** Rheobase measured from 5 ms-long current pulses. Mixed-effects model, sex effect F (1, 56) = 7.7, P = 0.007, ZT effect F (1, 56) = 1.4, P = 0.24, sex x ZT interaction F (1, 56) = 0.99, P = 0.32, n > 5 mice per sex, bars represent mean. **g**. Evoked activity in nociceptors between ZT2 and ZT14 in female and male mice. For females, two-way ANOVA with repeated measures, current effect F (2.4, 55) = 19, P < 0.001, ZT effect F (1, 23) = 0.00079, P = 0.98, current x ZT interaction F (8, 184) = 0.34, P = 0.95, n > 5 mice per group, mean ± SEM. For males, two-way ANOVA with repeated measures, current effect F (1.7, 57) = 34, P < 0.001, ZT effect F (1, 33) = 6.6, P = 0.015, current x ZT interaction F (8, 264) = 5.0, P < 0.001, n > 5 mice per group, mean ± SEM. **h.** Representation of the optogenetic experiment on Nav1.8-ChR2 mice. **i.** Paw withdrawal latency in Nav1.8-ChR2 female and male mice at ZT2 in comparison to ZT16. Two-way ANOVA with repeated measures and post-hoc Tukey tests, sex effect F (1, 7) = 1.9, P = 0.21, ZT effect F (1, 7) = 29, P = 0.001, sex x ZT interaction F (1, 7) = 21, P = 0.002, n > 4 mice per sex, bars represent mean. APs = action potentials, RMP = resting membrane potential. * P < 0.05, ** P < 0.01, *** P < 0.001.

### Evoked activity in nociceptors is reduced by the specific ablation of *Bmal1*

While we demonstrated the presence of an active molecular clock in the DRG, the daily fluctuations in nociceptor excitability may not be autonomously regulated by the nociceptors themselves but could instead be influenced by external factors; for instance, other cell types within the DRG could mediate these rhythmic changes through paracrine signaling, metabolic cues, or neuroimmune interactions ^26,27^. Additionally, systemic factors, such as hormonal fluctuations or signals from the central circadian clock in the suprachiasmatic nucleus, may also play a role in shaping the observed patterns of excitability ^28,29^. To test for autonomous regulation, we generated transgenic mice specifically lacking *Bmal1* expression in Nav1.8+ neurons ^30^ in order to specifically suppress the molecular clock in nociceptors with Bmal1^lox/lox^ and Nav1.8^cre/wt^;Bmal1^lox/lox^ littermates used as control and conditional knockout (cKO) mice, respectively. The efficiency of the cKO was assessed using RNA *in situ* hybridization, with significantly reduced expression of *Bmal1* at ZT2 (peak of *Bmal1* expression in WT mice) identified in cKO mice compared to control littermates (Extended Data Fig. 2a-b). Similar to our previous electrophysiology experiments, we recorded from nociceptors in whole-mount DRGs at ZT2 and ZT14 (Fig. 3a-b) to test if a nociceptor-autonomous clock can impact excitability. First, we observed an effect of the cKO on the resting membrane potential and rheobase of female nociceptors (Fig. 3a, c-d, Extended Data Fig. 2c-d), unlike male nociceptors (Fig. 3b, e-f, Extended Data Fig. 2e-f). This suggests a modification of the passive membrane properties in female mice lacking *Bmal1* in nociceptors. No effect of ZT was noticed, consistent with our results in wildtype C57BL/6J mice (Fig. 2e-f, Extended Data Fig. 2g-j). We next applied our step protocol. Surprisingly, while the evoked activity did not change over time in C57BL/6J females, the specific loss of *Bmal1* reduced evoked firing during the dark phase, but not in the light phase (Fig. 3g). In males, evoked firing at ZT2 was lower in cKO mice but was similar between the two genotypes at ZT14 (Fig. 3h). Overall, these findings suggest a cell-autonomous mechanism driven by the core circadian gene *Bmal1* that regulates the excitability of nociceptors in both male and female mice. Notably, ablation of the circadian machinery preferably impacts nociceptor activity in males in the light phase and in females in the dark phase.

**Fig. 3.**
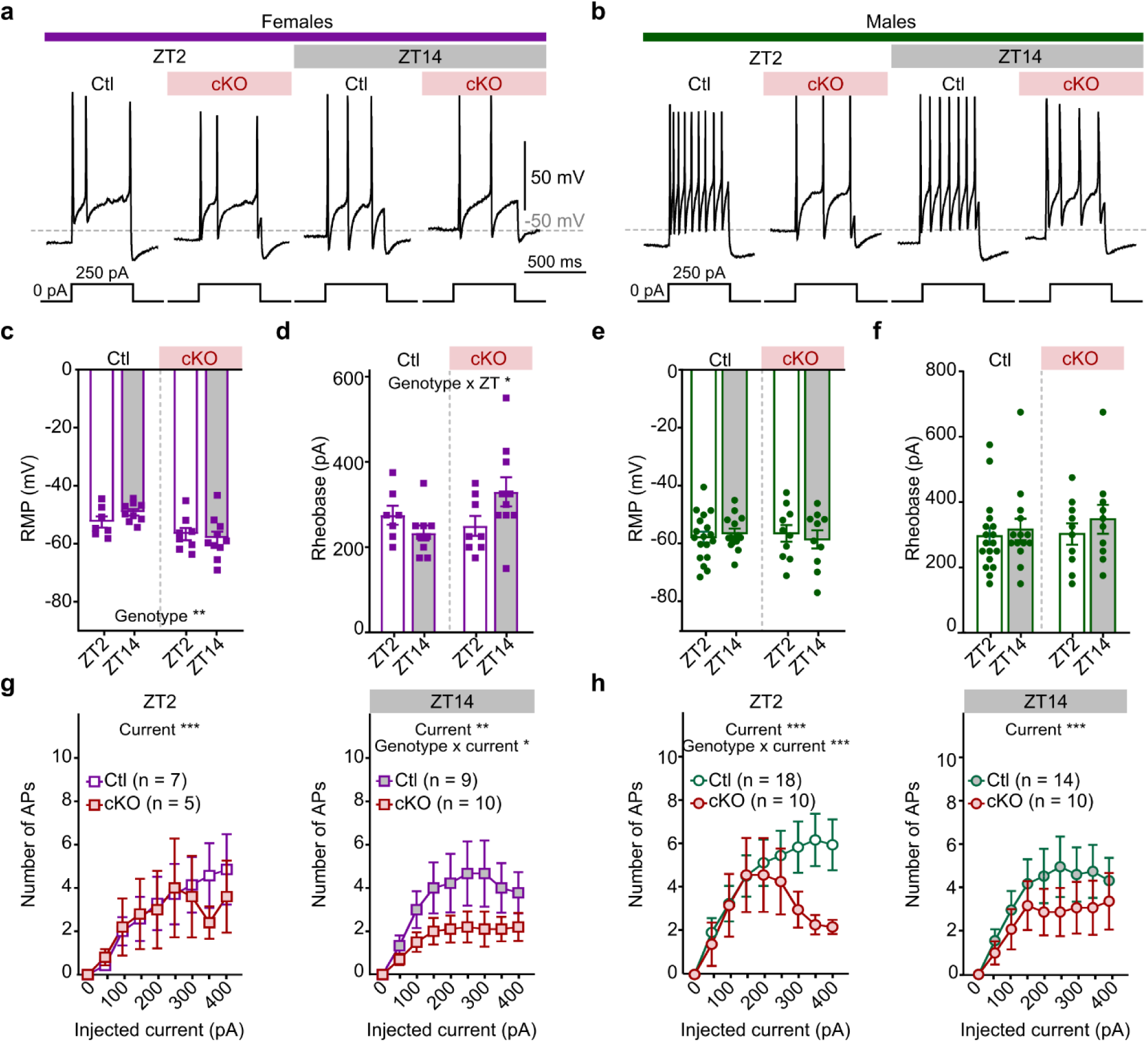
Specific *Bmal1* ablation in sensory neurons reduces excitability in both female and male mice. **a.** Evoked activity at 250 pA for 500 ms in female mice with *Bmal1* ablated in Nav1.8 neurons (cKO) and their control littermates (Ctl) at ZT2 and ZT14. **b.** Same as in a. for male mice. **c.** Resting membrane potential in Ctl and cKO female mice. Two-way ANOVA, genotype effect F (1, 30) = 11, P = 0.002, ZT effect F (1, 30) = 0.23, P = 0.63, genotype x ZT interaction F (1, 30) = 1.5, P = 0.24, n > 3 mice per genotype, bars represent mean. **d.** Rheobase in Ctl and cKO female mice. Two-way ANOVA, genotype effect F (1, 30) = 1.5, P = 0.23, ZT effect F (1, 30) = 0.26, P = 0.62, genotype x ZT interaction F (1, 30) = 5.4, P = 0.028, n > 3 mice per genotype, bars represent mean. **e.** Same as in c. for male mice. Two-way ANOVA, genotype effect F (1, 48) = 0.028, P = 0.87, ZT effect F (1, 48) = 0.022, P = 0.88, genotype x ZT interaction F (1, 48) = 0.56, P = 0.46, n > 3 mice per genotype, bars represent mean. **f.** Same as in d. for male mice. Two-way ANOVA, genotype effect F (1, 48) = 0.31, P = 0.58, ZT effect F (1, 48) = 0.92, P = 0.34, genotype x ZT interaction F (1, 48) = 0.13, P = 0.71, n > 3 mice per genotype, bars represent mean. **g.** Evoked activity at ZT2 (left) and ZT14 (right) in Ctl female mice compared to cKO. For ZT2, two-way ANOVA with repeated measures, current effect F (2, 20) = 7.3, P = 0.004, genotype effect F (1, 10) = 0.055, P = 0.82, current x genotype interaction F (8, 80) = 0.60, P = 0.78, n > 5 mice per group, mean ± SEM. For ZT14, two-way ANOVA with repeated measures, current effect F (1.9, 25) = 7.7, P = 0.003, genotype effect F (1, 13) = 3.0, P = 0.11, current x genotype interaction F (8, 104) = 2.1, P = 0.04, n > 5 mice per group, mean ± SEM. **h**. Same as in g. for male mice. For ZT2, two-way ANOVA with repeated measures, current effect F (2.3, 59) = 18, P < 0.001, genotype effect F (1, 26) = 1.1, P = 0.31, current x genotype interaction F (8, 208) = 4.4, P < 0.001, n > 5 mice per group, mean ± SEM. For ZT14, two-way ANOVA with repeated measures, current effect F (2, 44) = 12, P < 0.001, genotype effect F (1, 22) = 0.77, P = 0.40, current x genotype interaction F (8, 176) = 0.64, P = 0.74, n > 5 mice per group, mean ± SEM. APs = action potentials, RMP = resting membrane potential. * P < 0.05, ** P < 0.01, *** P < 0.001.

### Loss of *Bmal1* in nociceptors prevents changes in sensitivity to heat stimuli in male mice

Previous studies have demonstrated a correlation between the excitability of primary sensory neurons and nociception ^31,32^. We therefore performed behavioral tests on our genetically modified mice to confirm that our electrophysiological data are reflected at the behavioral level. First, we tested changes in the sensitivity of mice to a thermal stimulus using the Hargreaves radiant heat test (Fig. 4a). Withdrawal latency in female mice was mildly (post-hoc tests non-significant) impacted by the cKO and time of day (Fig. 4a). In males, we observed significantly increased withdrawal latency at ZT14 in control mice, corresponding to a diminished sensitivity to heat during the dark phase and matching the data obtained using the Nav1.8-ChR2 mice (Fig. 2h-i). This effect was lost in cKO male mice (Fig. 4a). These behavioral results match our electrophysiological recordings, with no change in excitability or heat sensitivity in female mice versus reduced nociceptor excitability and heat sensitivity in male mice during the dark phase. Sensitivity to cold and touch, assessed with the acetone test (Fig. 4b) and Von Frey test (Fig. 4b), respectively, was not impacted by daily rhythm in either control or cKO male mice. This suggests that cold and mechanical sensitivity involve clock-independent mechanisms in male mice. In contrast, ablation of the circadian clock in female nociceptors increased cold and mechanical sensitivity during the dark phase (Fig. 4b-c). Importantly, behavioral results obtained with Bmal1^lox/lox^ and Nav1.8^cre/wt^;Bmal1^lox/lox^ mice were replicated in Nav1.8^cre/wt^ and Nav1.8^cre/cre^;Bmal1^lox/lox^ mice, respectively (see Extended Data Fig. 3a-i).

**Fig. 4.**
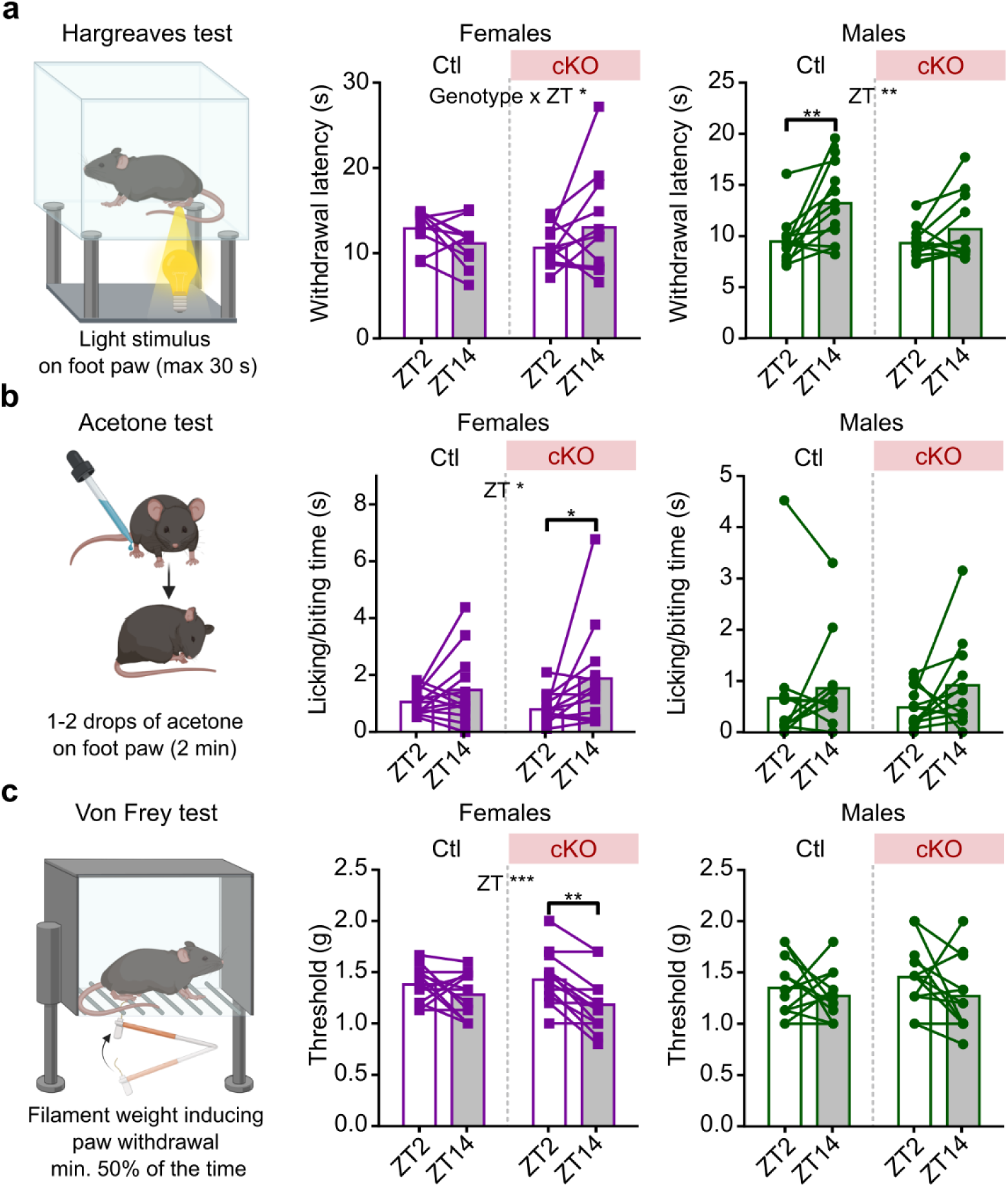
Specific *Bmal1* ablation in nociceptors affects sensitivity to different stimulus modalities. **a.** Hargreaves test (sensitivity to heat). Withdrawal latency at ZT2 and ZT14 in Ctl and cKO female and male mice. For females, two-way ANOVA with repeated measures and post-hoc Tukey tests, genotype effect F (1, 20) = 0.024, P = 0.88, ZT effect F (1, 20) = 0.11, P = 0.74, genotype x ZT interaction F (1, 20) = 4.6, P = 0.045, n > 10 mice per genotype, bars represent mean. For males, two-way ANOVA with repeated measures and post-hoc Tukey tests, genotype effect F (1, 22) = 2.0, P = 0.18, ZT effect F (1, 22) = 14, P = 0.001, genotype x ZT interaction F (1, 22) = 3.0, P = 0.096, n > 10 mice per genotype, bars represent mean. **b.** Acetone test (sensitivity to cold). Time spent licking/biting at ZT14 and ZT2 in Ctl and cKO female and male mice. For females, two-way ANOVA with repeated measures and post-hoc Tukey tests, genotype effect F (1, 24) = 0.048, P = 0.83, ZT effect F (1, 24) = 6.4, P = 0.019, genotype x ZT interaction F (1, 24) = 1.2, P = 0.28, n > 10 mice per genotype, bars represent mean. For males, two-way ANOVA with repeated measures and post-hoc Tukey tests, genotype effect F (1, 22) = 0.035, P = 0.85, ZT effect F (1, 22) = 3.0, P = 0.095, genotype x ZT interaction F (1, 22) = 0.42, P = 0.52, n > 10 mice per genotype, bars represent mean. **c.** Von Frey test (mechanical sensitivity). Mechanical threshold at ZT2 and ZT14 in Ctl and cKO female and male mice. For females, two-way ANOVA with repeated measures and post-hoc Tukey tests, genotype effect F (1, 24) = 0.091, P = 0.77, ZT effect F (1, 24) = 15, P < 0.001, genotype x ZT interaction F (1, 24) = 2.7, P = 0.11, n > 10 mice per genotype, bars represent mean. For males, two-way ANOVA with repeated measures and post-hoc Tukey tests, genotype effect F (1, 22) = 0.32, P = 0.58, ZT effect F (1, 22) = 2.3, P = 0.14, genotype x ZT interaction F (1, 22) = 0.39, P = 0.54, n > 10 mice per genotype, bars represent mean. * P < 0.05, ** P < 0.01, *** P < 0.001.

### Chloride currents impact the rhythms of male nociceptor excitability

Next, we aimed to identify the key molecular players regulated by the circadian clock that modulate fluctuations in nociceptor excitability in male mice and drive diurnal variations in heat pain sensitivity. By investigating clock-controlled genes and signaling pathways, we sought to determine how rhythmic changes in ion channel activity, neurotransmitter release, or intracellular signaling contribute to the daily modulation of pain sensitivity. We performed RNA sequencing on whole DRGs collected from C57BL/6J male mice at ZT2 and ZT14. Given that neuron excitability is intricately linked to transmembrane ion flux, we compared the transcriptomic changes in the expression of 295 genes involved in that process and expressed in our dataset (KEGG BRITE: brmmu04040). Candidate analysis revealed significantly increased expression of the chloride voltage-gated channel 2 (ClC-2, encoded by the *Clcn2* gene) at ZT14 relative to ZT2 (Fig. 5a); no other genes met our significance criteria (P_Bonferroni_ < 0.05). We next replicated these results using RT-qPCR on DRGs collected from male and female C57BL/6J mice and found a specific increase of *Clcn2* expression at ZT14 in males, while no changes in expression were observed in female mice across the day (Fig. 5b). The increase in *Clcn2* expression at ZT14 was lost in *Bmal1* cKO male mice (Fig. 5c), confirming the *Bmal1*-regulated expression of this channel, in a sex-specific manner, in nociceptors.

**Fig. 5.**
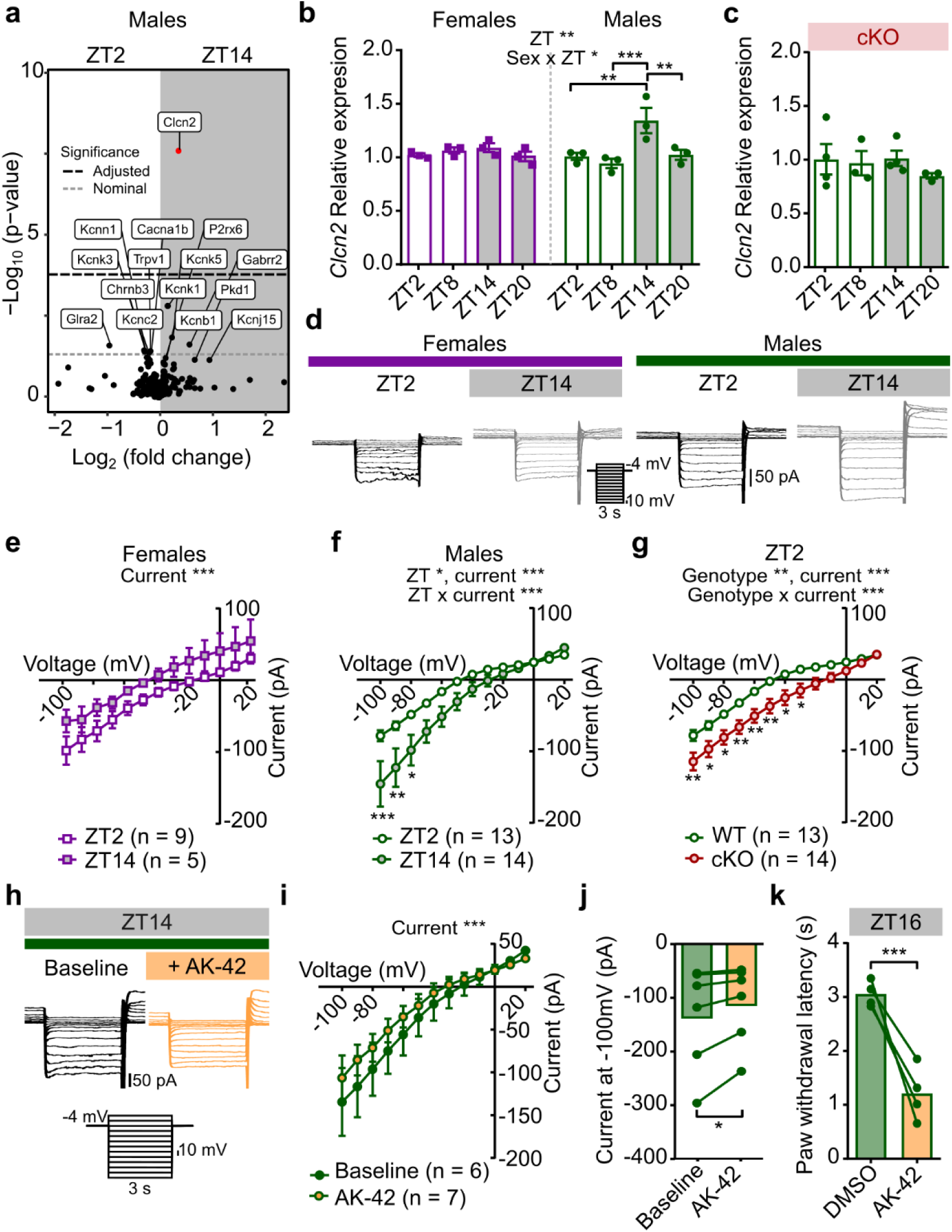
Voltage-gated chloride channels are more expressed and active at ZT14 in male mice. **a.** RNA seq candidate analysis of ion channel genes (KEGG br:mmu04040) in male mice DRG at ZT14 in comparison to ZT2 (n = 3 mice per time point; P_Bonferroni_ < 0.05). **b.** Relative expression of *Clcn2* measured by quantitative RT-PCR in female and male mice. Two-way ANOVA and post-hoc Tukey tests, sex effect F (1, 16) = 0.50, P = 0.49, ZT effect F (3, 16) = 7.1, P = 0.003, sex x ZT interaction F (3, 16) = 4.3, P = 0.022, n = 3 mice per group, mean ± SEM. **c.** Relative expression of *Clcn2* in cKO male mice. Kruskal-Wallis test, P = 0.40, n = 3 mice per group, mean ± SEM. **d**. Voltage clamp recordings of nociceptor chloride currents at ZT2 and ZT14 in WT female and male DRGs. **e.** Chloride currents at ZT2 and ZT14 in female nociceptors. Two-way ANOVA with repeated measures, current effect F (1.3, 15) = 28, P < 0.001, ZT effect F (1, 12) = 4.7, P = 0.051, current x ZT interaction F (12, 144) = 0.11, P > 0.99, n > 5 mice per group, mean ± SEM. **f.** Same as in e. for male mice. Two-way ANOVA with repeated measures and post-hoc Tukey tests, current effect F (1.2, 28) = 69, P < 0.001, ZT effect F (1, 25) = 4.5, P = 0.044, current x ZT interaction F (12, 300) = 4.3, P < 0.001, n > 5 mice per group, mean ± SEM. **g.** Chloride currents at ZT2 in male nociceptors from WT and cKO. Two-way ANOVA with repeated measures and post-hoc Tukey tests, current effect F (1.8, 45) = 168, P < 0.001, genotype effect F (1, 25) = 11, P = 0.003, current x genotype interaction F (12, 300) = 3.6, P < 0.001, n > 5 mice per group, mean ± SEM. **h.** Sample voltage clamp recording of a nociceptor at ZT14 from male before and after blocking ClC-2 with 1µM AK-42. **i.** Chloride currents recorded from male neurons before and after applying AK-42. Two-way ANOVA with repeated measures, current effect F (1.1, 13) = 45, P < 0.001, drug effect F (1, 11) = 0.38, P = 0.55, current x drug interaction F (12, 132) = 0.72, P = 0.73, n > 5 mice per group, mean ± SEM. **j.** Steady state current during −100 mV steps in neurons recorded before and after applying AK-42. Wilcoxon test, P = 0.031, n > 3 mice, bars represent means. **k**. Response latency of male Nav1.8-ChR2 mice to optogenetic stimulation after s.c. injection of 10µM AK-42 at ZT16 (n = 4 mice). Paired-t test, P = 0.009, n = 4 mice, bars represent means. * P < 0.05, ** P < 0.01, *** P < 0.001.

Because the ClC-2 channel is responsible for extruding chloride from the cell, we hypothesized that the increased expression of ClC-2 during the dark phase would increase chloride efflux under controlled conditions in nociceptors. To test this, we used voltage-clamp recordings to measure the chloride component of the leak conductance in our *ex-vivo* preparation. Briefly, chloride current was isolated by patching neurons with a cesium-based intracellular solution to block potassium channels and applying tetrodotoxin extracellularly to block sodium channels (see Methods). Current amplitude in female mice did not differ between ZT2 and ZT14 (Fig. 5d-e), consistent with the lack of detectable fluctuations for *Clcn2* in female miceIn males, on the other hand, the current amplitude at negative potential was higher at ZT14 than at ZT2, indicating greater efflux of chloride from the cell at night (Fig. 5d-f). Since ClC-2 channels are preferentially active at membrane potentials below resting potential, we hypothesized that the increase in inward current is due to increased membrane expression of ClC-2 channels during the dark phase. We then tested if deletion of *Bmal1* in nociceptors impacted chloride efflux specifically during the light phase, when an effect on excitability was observed at ZT2 in males. We identified a larger current in response to our step protocol in neurons from mice with disrupted clock machinery in nociceptors (Fig. 5g).

To test the role of ClC-2 in reducing nociceptor excitability in the dark phase, we bath applied AK-42, a recently developed specific potent inhibitor of ClC-2 ^33^, to nociceptors. Blocking ClC-2 significantly reduced chloride efflux at ZT14, and in so doing, abolished the difference in chloride efflux between ZT14 and ZT2 (Fig. 5h-j). Our electrophysiology work shows that nociceptors experience a functional change, that is mimicked by specific inhibition of ClC-2, which implies that these cells also experienced the transcriptional changes observed. Finally, to test if the increased levels of ClC-2 in nociceptors explains reduced nociception in males in the dark phase, we tested the response of male Nav1.8-ChR2 mice in the dark phase. Intraplantar injection of AK-42 significantly reduced the paw withdrawal latency to optogenetic stimulation (Fig. 5k), indicating that inhibition of ClC-2 in the dark phase is sufficient to restore a light-phase phenotype in male mice. Thus, rhythmic ClC-2 expression in nociceptors impacts both their excitability and basal nociception.

## DISCUSSION

Understanding the mechanisms controlling nociception in healthy organisms is necessary and fundamental to the development of efficient treatments for individuals suffering from acute and chronic pain. Recent studies have provided evidence that pain perception and modulation are dynamic processes influenced independently by circadian rhythms and sex differences. However, little is known about the potential interaction of these two factors, nor how the peripheral sensory nervous system could be implicated. In particular, nociceptors are of prime interest and serve as the initial relay in the pain pathway: they detect, encode, and transmit painful stimuli to the central nervous system ^34^. Variations in nociceptor physiology influence how nociceptive stimuli are perceived ^35^. Here we describe a rhythmic clock in nociceptors that modifies voltage-gated chloride channel expression in a sex-dependent manner, controlling nociceptor excitability and directly impacting pain-related behaviors.

Sex-specific differences in nociception are well-documented, with males typically exhibiting higher pain thresholds than females under normal conditions ^1,25^. However, studies paying attention to the interaction of sex and daily rhythm are rare. Most studies reporting diurnal fluctuation in pain sensitivity tested thermal stimuli on male rodents ^12,13,16,18,36^. Similar observations have also been carried out in humans, with males showing lower sensitivity to heat during their active (light) phase ^9^, matching our results in the active (dark) phase in male mice. While the ecological and/or evolutionary relevance of sexual dimorphism in the daily fluctuation of nociception in mice can be debated, it remains unclear if human females lack a circadian rhythm of pain as we find in mice. In this study, we suggest that the clock machinery in nociceptors is responsible for the sex-specific divergence of pain rhythmicity.

Whilst evidence indicates that genes expressed in human ^37^ and mouse DRGs ^37,38^ diverge between males and females, sex differences in clock machinery of nociceptors remains unexplored. Our study revealed some differences between male and female DRG clock genes, notably the absence of a rhythmic expression of *Nr1d2* in males and increased overall levels of *Per1* in females. Sex differences in the molecular clock have also been identified in peripheral tissues ^39–41^ and the brain ^42^. In DRGs, the presence of a functional molecular clock has been reported by former studies ^20–22^ but the sex-specificity of its expression patterns in DRG remained underexplored. Importantly, *Nr1d1/2* ^43–45^ and *Per1* ^46^ were recently identified as regulators of nociceptive hypersensitivity in mice. It is important to note, however, that our work was completed under light/dark cycle conditions; as such, environmental cues remain a confounder in our results and so our effects are determined to be diurnal and cannot be fully ascribed to circadian rhythms, which can only be assessed under specific conditions.

Ion channels play a key role in controlling the excitability of the central and peripheral neurons ^47^, and can show sex-specificity in the context of pain when assessed via electrophysiological methods across species ^29,48^. In particular, chloride currents have already been described as critical regulators of circadian rhythms ^49^. We identified ClC-2 as a potential key actor in the daily fluctuation of excitability and nociception in male mice. ClC-2 expression is ubiquitous, but previous research revealed its presence in neurons ^33,50,51^ and its particular role in regulating their excitability ^52,53^. Here, we highlight its diurnal (in animals kept in 12h light-dark cycles) and circadian control (in *Bmal1*-deficient mice) in a sex-dependent manner. These findings are in line with previous work showing that the transcription factors *Bmal1* and *Clock* can regulate *Clcn2* expression ^54^. While we show that inhibition of ClC-2 results in increased sensitivity during the dark phase, a previous study using the channel agonist lubiprostone showed reduced visceral hyperalgesia ^55^. Thus, targeting ClC-2 and/or its rhythmic expression in nociceptors may serve as a promising approach for therapeutic intervention in pain conditions.

Disruptions in circadian rhythms, such as altered light cycles or clock gene mutations, modify excitability and pain sensitivity. Dim light exposure or shifted light schedule results in altered nociceptive thresholds ^56,57^. Similarly, *Per2* mutant mice exhibit increased pain behaviors during the dark phase after formalin injection, correlated with elevated Substance P expression ^22^. Conditional *Bmal1* deletion in the hippocampus increased the excitability of neurons ^58^. In opposition to our finding, the specific loss of *Bmal1* in Nav1.8+ neurons has been shown to reduce osteoarthritis pain ^57^. This suggests that the effect of targeted *Bmal1* ablation on pain perception may vary depending on whether an individual is experiencing acute or chronic pain. Chronic ablation of the gene is also likely to cause compensatory changes that will depend on individual pain states. Loss of *Bmal1* in the healthy state may disrupt normal circadian regulation of nociceptor excitability, potentially altering pain sensitivity in a time-dependent manner. However, in the context of chronic pain, where nociceptive processing is already dysregulated, the consequences of *Bmal1* ablation can be more complex: it may either exacerbate pain by further disrupting homeostatic mechanisms or, conversely, reduce pain by interfering with maladaptive nociceptive plasticity. Further research is needed to determine how loss of *Bmal1* alters pain states and whether its effects are influenced by underlying inflammatory, neuropathic, or metabolic conditions.

Our study also revealed that disruption of the circadian clock in nociceptors directly impacted specific modalities of pain in a sex-dependent manner. Heat sensitivity was altered in male mice, whereas cold and touch sensitivity were altered in female mice. Most nociceptors respond to multiple stimulus modalities ^32,59,60^. Loss of Nav1.8 modifies the response to both thermal and mechanical stimuli in mice ^61,62^. However, only recently have sex differences in the polymodality of nociceptors been identified between males and females using electrophysiological recordings ^29^. This further suggests that the sexual dimorphism observed in our study is due in part to the intrinsic clock machinery and its downstream molecular cascade.

Surprisingly, unlike controls, female mice with disrupted expression of *Bmal1* in nociceptors had a higher pain sensitivity at night for these modalities. Since our electrophysiological data indicate reduced excitability at night in cKO female mice, we hypothesize that other sensory neuron subsets, satellite glia, or immune cells may affect these functional outcomes, independent of the intrinsic cell-autonomous excitability of female nociceptors ^17,56,63^. Immune cells and their secreted mediators, such as opioids, could act on nociceptors to reduce excitability to counterbalance the intrinsic, cell-autonomous properties of female nociceptors ^64,65^. In addition to immune modulation, recent findings have highlighted sexual dimorphism in the expression of prolactin and orexin receptors in nociceptors. Prolactin, a hormone involved in immune regulation and pain modulation, exhibits peak secretion during sleep, while orexin, a neuropeptide primarily associated with wakefulness and arousal, follows the opposite pattern ^66–68^. Given that prolactin and orexin levels are subject to circadian fluctuations, their rhythmic expression could further influence nociceptor excitability and pain perception across the day-night cycle. Further research is needed to determine how these factors could interact with the intrinsic circadian clock in nociceptors, and whether they act in a sexually-dimorphic manner to alter pain. Understanding these complex interactions could provide insight into sex-specific pain modulation and potential chronotherapeutic strategies for pain management.

Investigating the circadian rhythmicity of pain is often confounded by disrupted sleep, which can independently alter pain sensitivity ^7,69,70^. Moreover, most studies have focused on male subjects, leaving critical gaps in understanding sex-specific mechanisms. Our work addresses these limitations and disentangles the sex-based differences in the diurnal rhythmicity of pain, circadian gene expression in nociceptors, and sensory neuron excitability. Our convergent electrophysiological and behavioral results add support to our findings, and ultimately led to the identification of rhythmic ClC-2 expression in sensory neurons that specifically controls the sexually dimorphic rhythms of basal nociception.

## METHODS

### Animals

All work was performed in 6–12 weeks-old mice housed in light- and temperature-controlled rooms. Mice used in experiments include C57BL/6J (Jackson Labs, stock #000664), B6.129-Scn10a<tm2(cre)Jnw/H (Nav1.8-Cre, kindly provided by Dr. John Wood [University College London, UK]), and B6.129S4(Cg)-*Bmal1*tm1Weit/J (*Bmal1*^Lox^, Jackson Labs, stock #007668). Mice that express ChR2-eYFP in nociceptive DRG neurons were generated by crossing homozygous Ai32 mice (Jackson Labs, stock #012569), which express ChR2(H134R)-eYFP in the presence of Cre recombinase, with Nav1.8-Cre mice (Tg(Scn10a-cre)1Rkun) kindly provided by Rohini Kuner (Heidelberg University, Germany). All mice were conventionally housed on a normal 12h/12h light/dark cycle except Nav1.8-ChR2 mice which were housed in a 14/10 environment, with food and water available *ad libitum* in the animal care facilities of Queen’s University (all strains except the Nav1.8-ChR2 line) and the Hospital for Sick Children. All procedures were approved by the Animal Care Committee at Queen’s University (#2019–1963) and at The Hospital for Sick Children (#65769) and were conducted in accordance with guidelines from the Canadian Council on Animal Care. Mice were euthanized with an intraperitoneal injection of sodium pentobarbital (200 mg/kg) followed by decapitation at the indicated ZT, with ZT0 corresponding to the light-onset. Appropriate statistical tests were performed using Prism software (GraphPad, La Jolla, CA).

### Pain behavior assays

Mice were habituated for 1h daily in individual compartments for each behavior assay. Mechanical threshold was measured using von Frey monofilaments (Ugo Basile) and defined as the minimum filament weight needed to elicit at least five responses (fast paw withdrawal, flinching, or licking/biting of the stimulated paw) over a total of 10 stimulations to determine the 50% threshold. For the cold hypersensitivity assessment, a drop of acetone was applied to the plantar area of the left hind paw and the time the mouse spent licking and biting the paw was recorded. The duration of the reaction was measured and analyzed as cumulative reaction time. The Hargreaves Radiant Heat Test (IITC Life Science, Woodland Hills, CA) was used to assess paw withdrawal latency in response to heat stimulation, according to the Hargreaves method^71^. Animals were first acclimated in plastic observation boxes for 20 min. Then, the mid-plantar surface of the mouse hind paw was exposed to a radiant heat source through a glass floor until paw withdrawal. The heat intensity was adjusted to produce a baseline of about 15 seconds in naïve mice, with a maximum cutoff of 30 s. Each mouse was tested three times, with at least two minutes between recordings. The average of all recordings was used for statistical analysis. Three baseline measurements were then taken on separate days for mechanical threshold and thermal latency to response and averaged. All behavioral experiments were carried out using at least two independent cohorts of mice.

### Automated behavior assays

RAMalgo was used to apply optogenetic stimuli transcutaneously to the paw ^72^. For the experiments, mice were housed with a 14 h/10 h light/dark cycle. Mice expressing ChR2 in nociceptors were transported from the housing room to the testing room for 1 hour of acclimation at ZT2 and ZT16 for several days before testing. On testing days, mice were acclimated for 1 hour before starting tests. At acclimation and testing at ZT16, cages were handled under red light and covered during transport to avoid exposing the mice to light. Testing at ZT16 was performed in the dark whereas testing at ZT2 was performed under regular lighting. Mice were kept in individual cubicles on a plexiglass platform illuminated with infrared light and viewed from below with an infrared-sensitive camera. All testing was performed with the experimenter controlling RAMalgo remotely from outside the room. Each mouse was tested with 3 trials per session with at least a 5-min interval between trials. For optogenetic stimulation, blue light (455 nm) was focused to a spot 3.5 mm in diameter centered on the paw, and its intensity ramped from 0 to 0.9 mW/mm2 (0-4 V command signal) over 15 seconds ^73^. Light intensity on target was measured with a s170C photodiode and PM100D optical power meter (Thorlabs). Photostimulation stopped automatically when the paw was withdrawn or at the 15-s cutoff. Paw withdrawal was measured automatically with millisecond precision by RAMalgo using red light reflected off the paw. AK-42 (10 μM, 20 μl) or the same volume of vehicle (DMSO diluted in saline) was injected subcutaneously (s.c) into the left hindpaw 30 min prior to testing. AK-42 was a gift from Justin Du Bois (Stanford University, USA) and was first diluted in Dimethyl sulfoxide (DMSO; BP231-1, ThermoFisher Scientific) at 0.1 M.

### RNA in situ hybridization

DRGs from Nav1.8-*Bmal1* KO and their littermate control were dissected, post-fixed in 4% Paraformaldehyde/PBS (Sigma–Aldrich, St. Louis, MO) for 2 hr, cryoprotected in 30% sucrose/PBS at 4°C, embedded in Pelco® cryo-embedding compound (Ted Pella, Redding, CA, USA, cat.27300), and frozen at −80℃. A Leica CM1950 cryostat (Leica, Wetzlar, Germany) was used to cut 14 μm cryosections were cut onto Superfrost™ Plus slides (Fisher Scientific, Waltham, MA, USA, cat.12-550-15). The RNAscope™ Multiplex Fluorescent V2 Assay was used for in situ hybridization (ACDBio, Newark, CA, USA). Manufacturer’s protocol was used for RNA in situ hybridization. Briefly, sections were washed with PBS to remove cryo-embedding compound, baked at 60℃ for 30 minutes, and post-fixed in 4% PFA for 15 minutes at 4℃. Tissue was dehydrated using an ethanol gradient with 50%, 70%, and 100% ethanol. Sections were incubated with hydrogen peroxide for 10 minutes, then washed with water.

Slides were incubated in target retrieval reagent heated up to at least 99℃ in a steamer (Hamilton Beach, Glen Allen, VA, USA, cat. 37530C) for 15 minutes. Sections were then dried and an ImmEdge™ hydrophobic barrier pen (Vector Laboratories, Newark, CA, USA, cat. H-4000) was used to draw a barrier around the tissue. Sections were incubated with RNAscope Protease III for 30 minutes at 40℃. Probes for Avil (ACDBio, cat. 498531), Scn10a (ACDBio, cat. 426011-C2), and Arntl (cat. 438741-C3) were used to confirm ablation of BMAL1 in Nav1.8 positive neurons. Probe mixture was added to slides and incubated for 2 hours at 40℃. Slides were then washed with wash buffer. Slides were incubated with three amplification solutions: Amp1, Amp2, and Amp3, consecutively, with Amp1 and Amp2 incubations lasting 30 minutes, and Amp3 incubation lasting 15 minutes, all at 40℃, with washes in between incubations. Next HRP-C1 signal was developed (horseradish peroxidase, channel 1) by incubating tissue in HRP-C1 reagent for 15 minutes at 40℃. Slides were then washed, then incubated with Opal™ 520 fluorophore (1:750 dilution) (Akoya Biosciences, Marlborough, MA, USA, cat. FP1487001KT) for 30 minutes at 40℃. Then the slides were washed at HRP blocker was added to the slides and incubated for 15 minutes at 40℃. The same incubation conditions and times were used to develop HRP-C2 (channel 2) and HRP-C3 (channel 3) signals, consecutively. For C2, Opal™ 570 fluorophore was used at 1:750 dilution (Akoya Biosciences, Marlborough, MA, USA, cat.FP1488001KT), and for C3 Opal™ 690 fluorophore was used at 1:750 dilution (Akoya Biosciences, Marlborough, MA, USA, cat. FP1497001KT). Fluoroshield™ mounting media with DAPI was then added to the slides and cover glass was applied (Fisher Scientific, Waltham, MA, USA, cat.22-050-228).

A Nikon AX/AX R confocal microscope was used to image hybridized DRG. For quantification, at least eight distinct 10× fields of view of lumbar DRG staining from n = 3 animals for each sex and genotype, were analyzed for co-localization and neuronal proportions. No sex difference was observed in conditional knockout efficacy, so males and females of the same genotype were pooled for statistical analysis. Statistical analysis and graphs were generated using Prism software (GraphPad, La Jolla, CA).

### RNA extraction

DRG tissues (L1 - L6) were dissected and stored at −80°C after mouse euthanasia at either ZT2, 8, 14, or 20. RNA was extracted using the RNeasy mini kit (Qiagen 74004). Pelleted and washed RNA was resuspended in RNase-free water and quantified using the NanoDrop system (Thermo Fisher, Waltham, MA). The concentration of each total RNA sample was standardized to 50 ng/µl. For RNA sequencing, three mice were pooled for each sample, and there were three samples for each time point.

### RNA sequencing

After tissues were harvested, they were flash-frozen and transferred to the Beijing Genomics Institute (BGI, Inc. One Broadway, 14th Floor, Cambridge, MA 02142 USA) for RNA extraction and sequencing with the DNBSEQ platform. The resulting sequencing reads were 150 base pair paired-end reads. BGI used SOAPnuke v1.5.2^74^ with the parameters “-l 15 -q 0.2 -n 0.05” to clean raw-sequencing reads. Data were shipped via physical hard drive and stored by the Queen’s University Centre for Advanced Computing.

Our curated workflow for transcriptomics data analysis is summarized in Extended Data Fig. 4a and detailed in the following sections. We evaluated the quality of sequencing reads with FastQC v0.11.9^75^ and MultiQC v1.12 ^76^. We filtered for base quality with Trimmomatic v0.36 ^77^ and aligned reads to the mouse genome (GENCODE release M24) with Hisat2 v2.2.1 ^78,79^. To evaluate alignment performance, we calculated the overall and unique alignment rates; all rates were > 90% (Extended Data Fig. 4b). Gene expression was quantified with StringTie v2.1.5, R v4.2.1^80^, and IsoformSwitchAnalyzeR v1.17.04 ^81,82^.

Downstream analyses were performed in R v3.6.0^83^. We performed outlier detection and median ratios normalization using the R packages arrayQualityMetrics v3.42.0 and DESeq2 v1.26.0 ^84,85^. arrayQualityMetrics uses three metrics to consider a sample an outlier: 1) its sum of distances to other samples, 2) the Kolmogorov-Smirnov statistic, and 3) the Hoeffding’s D-statistic. Samples would be removed if marked an outlier before and after normalization, or if multiple metrics marked them an outlier after normalization. No samples were removed. We used principal component analysis on variance stabilizing transformed counts to visualize the clustering of samples with DESeq2 (Extended Data Fig. 4c). Finally, we considered genes to be expressed if their median absolute deviation of normalized counts were >= 25% (0.015571). There were 29,206 genes remaining for downstream analyses.

To assess whether ion channel genes are different ZT2 versus ZT14 at the gene-level in the DRG, differential expression analyses of candidate genes were performed using DESeq2. Differentially expressed genes (DEGs) had a Bonferroni adjusted p-value < 0.05. Genes from the KEGG BRITE hierarchy of *Mus musculus* ion channels (br:mmu04040) were tested.

### Quantitative real-time PCR

Complementary DNA (cDNA) was produced using the High-Capacity cDNA Reverse Transcription Kit (Applied Biosystems™ 4368813). The PCR amplifications were performed in duplicate at 50°C for 120 s then 95°C for 20 s, followed by 40 cycles of thermal cycling at 95°C for 1 s and 60°C for 20 s on an Applied Biosystems 7500 machine (Life Technologies) using TaqMan™ Fast Advanced Master Mix (Applied Biosystems™ 4444558). Gapdh was used as an endogenous control to normalize differences. Quantification was performed by normalizing Ct (cycle threshold) values with Gapdh Ct and analyzed with the 2- ΔΔcT method. Gene expression was analyzed by using QuantStudio 7 Flex Real-Time PCR. The primers used in this study were: *Arntl/Bmal1* probe Mm00500226_m1, *Clock* probe Mm00455950_m1, *Nr1d1* probe Mm00520708_m1, *Nr1d2* probe Mm01310356_g1, *Per1* probe Mm00501813_m1, *Per2* probe Mm00478099_m1, *Cry1* probe Mm00514392_m1, *Cry2* probe Mm01331539_m1, *Clcn2* probe Mm01344255_g1 (ThermoFisher, Waltham, MA).

### Whole-cell recordings

At ZT2 or ZT14, DRGs (L3 or L4) were quickly removed from the vertebral column and placed in oxygenated aCSF containing (in mM): NaCl (123), NaHCO3 (26), KCl (3), NaH2PO4 (1.25), MgCl2 (1), CaCl2 (2), Glucose (11) adjusted to pH 7.4 with NaOH and osmolarity to 300 mOsm. Ventral root, connective tissue and the epineurium were carefully removed under a microscope. One ganglion was then transferred to the recording chamber of an upright fixed-stage BX51WI microscope (Olympus Corporation) equipped with differential interface contrast (DIC) optics at 40X objective magnification through a CCD camera. The intact DRG was continuously perfused (1 mL/min) with 95 % O2 and 5 % CO2-saturated aCSF at room temperature. A small area of the surface of the DRG was then digested with a glass patch pipette (∼1 MΩ) filled with 13 U/mL Collagenase (Liberase TM) for 15 min. Only small cells (< 40 pF and < 25 µm) were chosen for our recordings. A multiclamp 700B amplifier (Molecular Devices), an InstruTECH ITC-18 digitizer (HEKA Instruments Inc) and the Axograph software (Axograph Scientific) were used for recordings. Data were sampled at 5 kHz, Bessel filtered at 10 kHz and 1 kHz for current clamp and voltage clamp recordings respectively. Cell capacitance transients were automatically canceled by the capacitive cancellation circuitry on the amplifier. Leak junction potential was automatically compensated (∼10 mV). Patch clamp pipettes were pulled from 1.5 mm thin wall borosilicate glass capillaries (World Precision Instrument) on a P-97 (Sutter Instruments) and had a resistance between 5-10 MΩ, when filled with the pipette solution. All recordings were performed for up to 5 hours after mice euthanasia.

For current-clamp recordings, the intracellular solution used contained (in mM): K Gluconate (130), KCl (10), HEPES (10), EGTA (10), MgCl2 (2), MgATP (2), NaGTP (0.5) adjusted to pH 7.4 with KOH and osmolarity to 280 mOsm. After obtaining whole-cell configuration, cells were held at −60 mV for at least 5 min to allow equilibrium. RMP was obtained by averaging 1 minute of recording. A series of 5 ms current pulses (25 pA increment) were injected into the neuron to measure rheobase, i.e. the weakest pulse intensity able to evoke an action potential. Evoked activity was measured with 500 ms-long current steps tested in 25-50 pA increments. Data were discarded if: RMP was more depolarized than −40 mV; AP amplitude was < 50 mV; input resistance was < 200 MΩ; series resistance was > 40 MΩ; or series and membrane resistance changed by > 25 % during the recording. AP amplitude was defined as the peak of the AP from the RMP.

For voltage-clamp experiments, the intracellular solution used contained (in mM): CsCl (130), HEPES (10), EGTA (10), MgATP (3), Na2GTP (0.3); and the bath contained: NaCl (116), NaHCO3 (26), CsCl (10), KCl (2.5), sodium phosphate (1.25), MgSO4 (1), glucose (10), EGTA (2), tetrodotoxin (1) (TTX-550, Alomone Labs). After obtaining whole-cell configuration, cells were held at −4 mV. Series of 3 s steps from −100 mV to +20 mV were applied every 30 s to produce the IV curve. Data were discarded if: an average inward current < −50 pA was measured at 0 mV; current measured before the steps varied by more than 100 pA; input resistance was < 200 MΩ; series resistance was > 40 MΩ; or series resistance changed by > 40 % during the recording. For AK-42 experiments, cells were recorded before (baseline) and 5 min after AK-42 was added to the bath.

## Acknowledgements

This work was funded by grants from the Canadian Institutes of Health Research (PJT-497592 to NG and Foundation Grant 167276 to SAP), Natural Sciences and Engineering Research Council of Canada (RGPIN-05604 to NG), and MS Canada (grant #3316 to NG). AB was funded by a postdoctoral fellowship from MS Canada. The authors thank Dr. Merritt Maduke (Dept of Molecular and Cellular Physiology, Stanford University) for advice regarding the use of AK-42. Computations were performed on resources and with support provided by the Centre for Advanced Computing (CAC) at Queen’s University in Kingston, Ontario. The CAC is supported in part by funding from Queen’s University, the Digital Research Alliance of Canada and the Government of Ontario.

## Author contributions

AB and NG conceived the project; AB, SAP, QD, and NG designed the project; AB, CAB, YFX, CD, CDO, LLB, and SDM performed experiments; AB, AMZ, YFX, CD, CDO, QD, SAP, and NG analyzed data; AB and NG wrote the manuscript with comments from all authors.

## Materials & Correspondence

All correspondence to Nader Ghasemlou.

## Code availability

Upon publication, scripts used to analyze data and generate figures will be available at https://github.com/ComputationalGenomicsLaboratory/naiveDRG and https://github.com/AurelieBre/DRGPatchClampProcessing.

## Data availability

Raw RNA sequencing data generated in the present study will be available in the National Center for Biotechnology Information’s Gene Expression Omnibus database repository.

## SUPPLEMENTARY DATA

**Extended Data Fig. 1.**
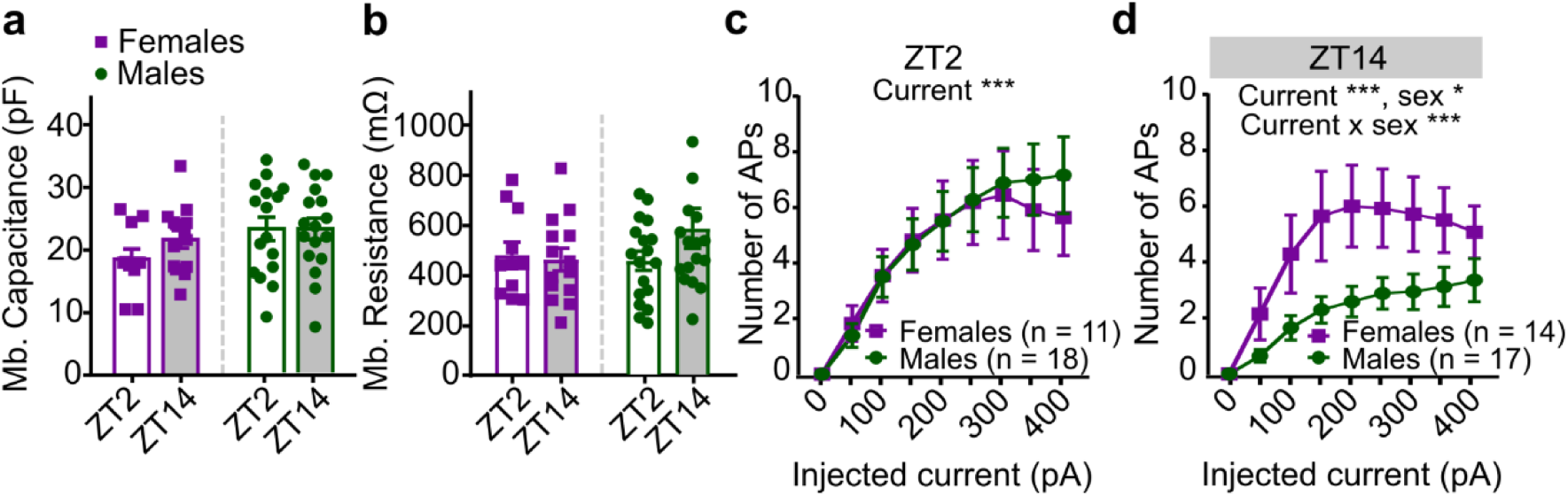
**a.** Membrane capacitance from recorded DRG neurons according time points and sex. Two-way ANOVA, sex effect F (1, 54) = 2.8, P = 0.100, ZT effect F (1, 54) = 0.84, P = 0.36, sex x ZT interaction F (1, 54) = 0.86, P = 0.36, n > 5 mice per group, mean ± SEM. **b.** Membrane resistance from recorded DRG neurons according time points and sex. Two-way ANOVA, sex effect F (1, 56) = 0.55, P = 0.46, ZT effect F (1, 56) = 0.87, P = 0.35, sex x ZT interaction F (1, 56) = 1.6, P = 0.22, n > 5 mice per group, mean ± SEM. **c.** Evoked action potential induced by a step protocol of 500 ms at ZT2 for male and female DRG. Two-way ANOVA with repeated measures, current effect F (1.7, 47) = 372 P < 0. 001, sex effect F (1, 27) = 0.039, P = 0.845, current x sex interaction F (8, 216) = 0.57, P = 0.80, n > 5 mice per group, mean ± SEM. **d.** Evoked action potential induced by a step protocol of 500 ms at ZT14 for male and female DRG. Two-way ANOVA with repeated measures and post-hoc Tukey tests, current effect F (2.3, 67) = 18, P < 0.001, sex effect F (1, 29) = 5.0, P = 0.034, current x sex interaction F (8, 232) = 2.1, P < 0.034, n > 5 mice per group, mean ± SEM. APs = action potentials. * P < 0.05, ** P < 0.01, *** P < 0.001.

**Extended Data Fig. 2.**
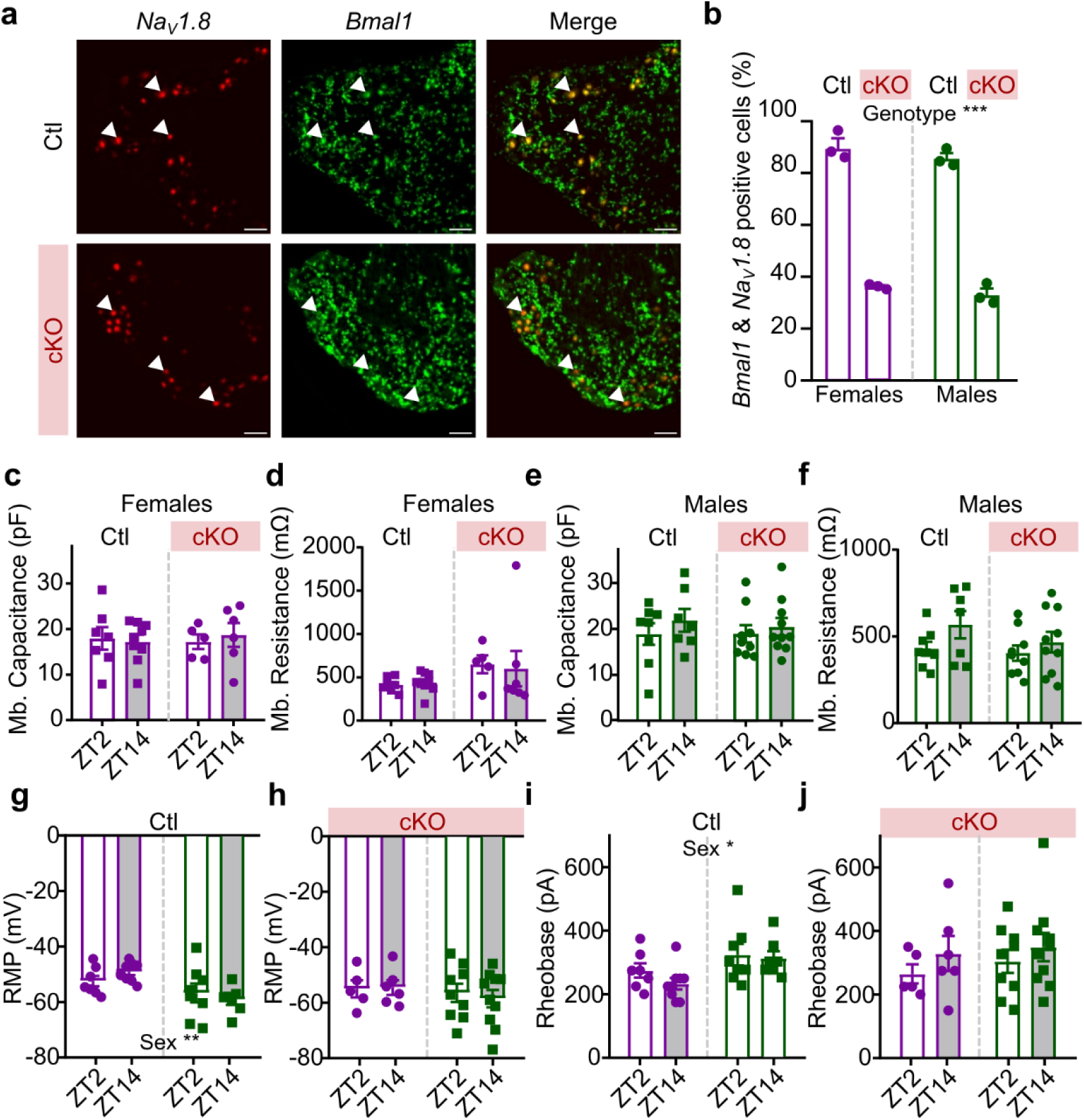
**a.** Co-staining of Bmal1 and Nav1.8 in L3/4 DRGs of Ctl (top) and cKO (bottom) male mice at ZT2 using RNA in situ hybridization. **b.** Statistical analysis using two-tailed unpaired t test. Two-way ANOVA, sex effect F (1, 8) = 1.9, P = 0.201, genotype effect F (1, 8) = 595, P < 0.001, sex x genotype interaction F (1, 8) = 0.11, P = 0.74, n = 3 mice per group, mean ± SEM. **c.** Membrane capacitance from recorded DRG neurons of cKO and Ctl female mice. Two-way ANOVA, ZT effect F (1, 23) = 0.036, P = 0.85, genotype effect F (1, 23) = 0.033, P = 0.86, ZT x genotype interaction F (1, 23) = 0.27, P = 0.61, n > 5 mice per group, mean ± SEM. **d.** Membrane resistance from recorded DRG neurons of cKO and Ctl female mice. Two-way ANOVA, ZT effect F (1, 24) = 0.0097, P = 0.92, genotype effect F (1, 24) = 3.0, P = 0.097, ZT x genotype interaction F (1, 24) = 0.11, P = 0.74, n > 5 mice per group, mean ± SEM. **e**. Same as in c. for male mice. Two-way ANOVA, ZT effect F (1, 30) = 1.1, P = 0.30, genotype effect F (1, 30) = 0.10, P = 0.75, ZT x genotype interaction F (1, 30) = 0.12, P = 0.73, n > 5 mice per group, mean ± SEM. **f.** Same as in d. for male mice. Two-way ANOVA, ZT effect F (1, 30) = 3.1, P = 0.086, genotype effect F (1, 30) = 1.2, P = 0.27, ZT x genotype interaction F (1, 30) = 0.44, P = 0.51, n > 5 mice per group, mean ± SEM. **g.** Resting membrane potential of Ctl female and male mice at ZT2 and ZT14. Two-way ANOVA, ZT effect F (1, 27) = 0.038, P = 0.85, sex effect F (1, 27) = 11, P = 0.003, ZT x sex interaction F (1, 27) = 1.7, P = 0.20, n > 5 mice per group, mean ± SEM. **h.** Resting membrane potential of cKO female and male mice at ZT2 and ZT14. Two-way ANOVA, ZT effect F (1, 26) = 0.050, P = 0.82, sex effect F (1, 26) = 0.74, P = 0.40, ZT x sex interaction F (1, 26) = 0.14, P = 0.714, n > 5 mice per group, mean ± SEM. **i.** Rheobase of Ctl female and male mice at ZT2 and ZT14. Two-way ANOVA, ZT effect F (1, 27) = 1.1, P = 0.30, sex effect F (1, 27) = 6.2, P = 0.019, ZT x sex interaction F (1, 27) = 0.37, P = 0.55, n > 5 mice per group, mean ± SEM. **j.** Rheobase of cKO female and male mice at ZT2 and ZT14. Two-way ANOVA, ZT effect F (1, 26) = 1.4, P = 0.25, sex effect F (1, 26) = 0.37, P = 0.55, ZT x sex interaction F (1, 26) = 0.044, P = 0.83, n > 5 mice per group, mean ± SEM. APs = action potentials, Mb. = membrane, RMP = resting membrane potential. * P < 0.05, ** P < 0.01, *** P < 0.001.

**Extended Data Fig. 3.**
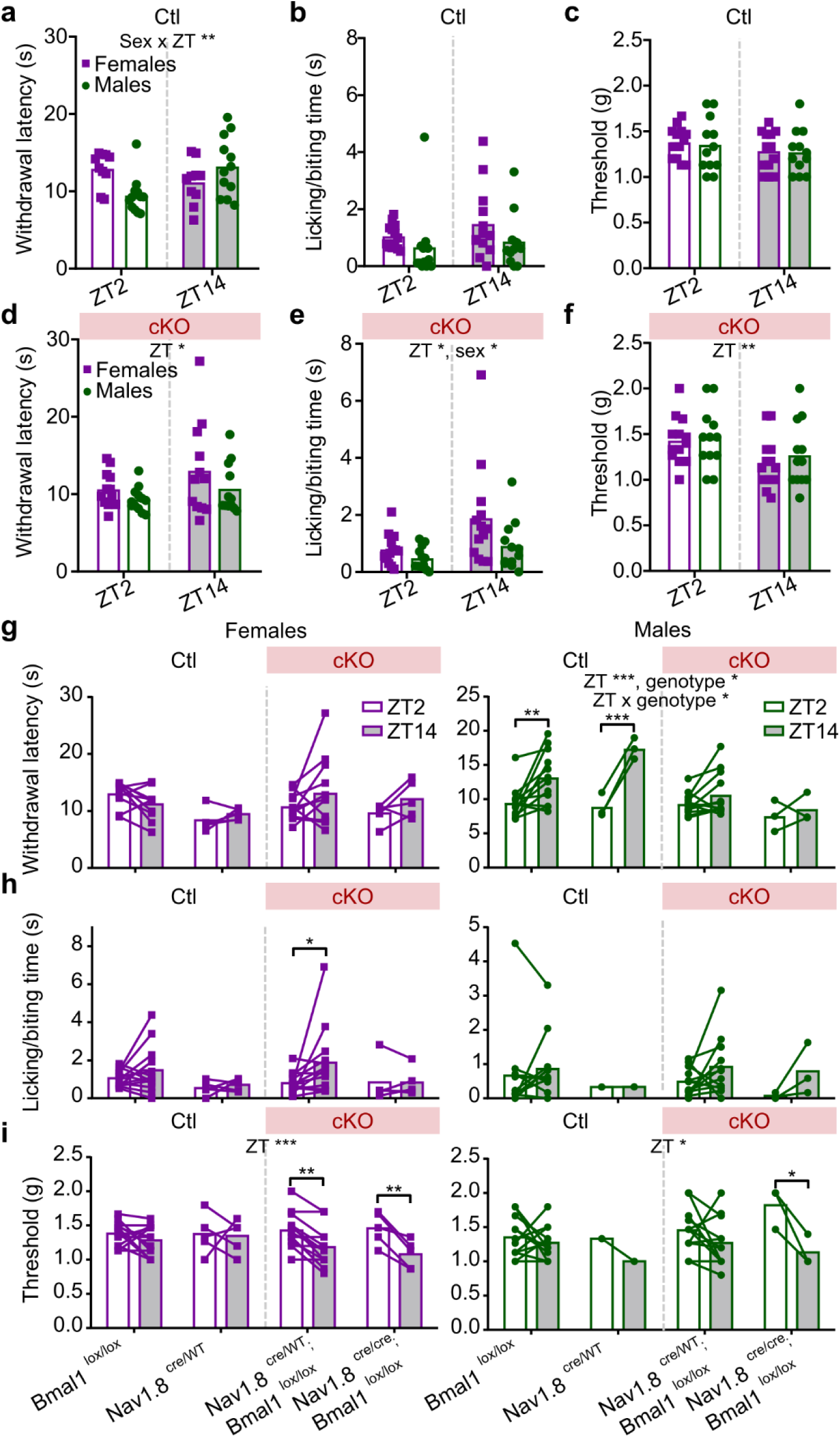
**a.** Withdrawal latency of Ctl female and male mice at ZT2 and ZT14. Two-way ANOVA, ZT effect F (1, 20) = 1.8, P = 0.20, sex effect F (1, 20) = 0.50, P = 0.49, ZT x sex interaction F (1, 20) = 13, P = 0.002, n > 5 mice per group, mean ± SEM. **b.** Licking/biting time of Ctl female and male mice at ZT2 and ZT14. Two-way ANOVA, ZT effect F (1, 23) = 2.4, P = 0.14, sex effect F (1, 23) = 2.0, P = 0.173, ZT x sex interaction F (1, 23) = 0.31, P = 0.58, n > 5 mice per group, mean ± SEM. **c.** Filament threshold of Ctl female and male mice at ZT2 and ZT14. Two-way ANOVA, ZT effect F (1, 23) = 1.7, P = 0.21, sex effect F (1, 23) = 0.095, P = 0.76, ZT x sex interaction F (1, 23) = 0.021, P = 0.89, n > 5 mice per group, mean ± SEM. **d.** Same as in a. for cKO mice. Two-way ANOVA, ZT effect F (1, 22) = 4.5, P = 0.045, sex effect F (1, 22) = 2.3, P = 0.14, ZT x sex interaction F (1, 22) = 0.34, P = 0.56, n > 5 mice per group, mean ± SEM. **e.** Same as in b. for cKO mice. Two-way ANOVA, ZT effect F (1, 23) = 6.7, P = 0.017, sex effect F (1, 23) = 4.3, P = 0.050, ZT x sex interaction F (1, 23) = 1.2, P = 0.28, n > 5 mice per group, mean ± SEM. **f.** Same as in c. for cKO mice. Two-way ANOVA, ZT effect F (1, 23) = 10, P = 0.004, sex effect F (1, 23) = 0.30, P = 0.59, ZT x sex interaction F (1, 23) = 0.19, P = 0.67, n > 5 mice per group, mean ± SEM. **g.** Comparison of two different Ctl transgenic mice lines (Bmal1^lox/lox^ and Nav1.8^cre/wt^) and two cKO mice lines (Nav1.8^cre/wt^;Bmal1^lox/lox^ and Nav1.8^cre/cre^;Bmal1^lox/lox^) at the Hargreaves test for both females (left) and males (right) at ZT2 and ZT14. For females, mixed-model effects and post-hoc Tukey tests, ZT effect F (1, 24) = 1.3, P = 0.26, genotype effect F (1.2, 13) = 2.3, P = 0.15, ZT x genotype interaction F (3, 32) = 1.6, P = 0.21, n > 5 mice per group, bars represent means. For males, two-way ANOVA and post-hoc Tukey tests, ZT effect F (1, 26) = 25, P < 0.001, genotype effect F (3, 26) = 3.3, P = 0.036, ZT x genotype interaction F (3, 26) = 4.5, P = 0.011, n > 5 mice per group, bars represent means. **h.** Same as in g. for acetone test. For females, mixed-model effects and post-hoc Tukey tests, ZT effect F (1, 32) = 2.8, P = 0.10, genotype effect F (3, 32) = 1.3, P = 0.29, ZT x genotype interaction F (3, 32) = 1.1, P = 0.346, n > 5 mice per group, bars represent means. For males, two-way ANOVA and post-hoc Tukey tests, ZT effect F (1, 24) = 1.6, P = 0.22, genotype effect F (3, 24) = 0.21, P = 0.88, ZT x genotype interaction F (3, 24) = 0.39, P = 0.76, n > 5 mice per group, bars represent means. **i.** Same as in g. for Von Frey test. For females, mixed-model effects and post-hoc Tukey tests, ZT effect F (1, 32) = 16, P < 0.001, genotype effect F (3, 32) = 0.20, P = 0. 90, ZT x genotype interaction F (3, 32) = 2.5, P = 0.081, n > 5 mice per group, bars represent means. For males, two-way ANOVA and post-hoc Tukey tests, ZT effect F (1, 24) = 6.4, P = 0.018, genotype effect F (3, 24) = 0.68, P = 0.57, ZT x genotype interaction F (3, 24) = 1.8, P = 0.18, n > 5 mice per group, bars represent means. * P < 0.05, ** P < 0.01, *** P < 0.001.

**Extended Data Fig. 4.**
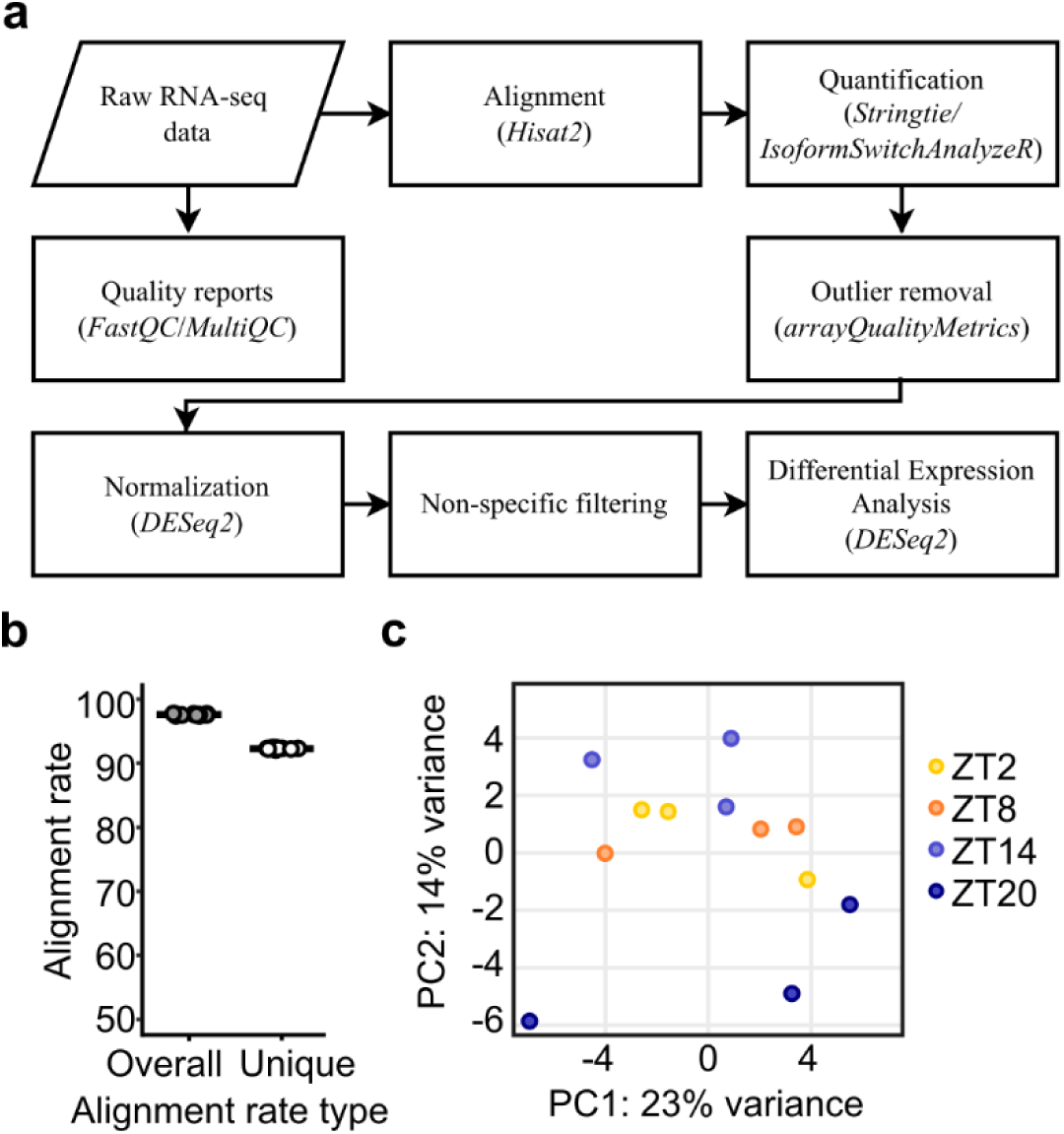
**a.** Curated workflow for transcriptomics data analysis. **b.** Overall and unique alignment rates. **c**. Principal component analysis on variance with *DESeq2*.

